# Intranasal oxytocin modulates very early visual processing of emotional faces

**DOI:** 10.1101/2021.04.15.440078

**Authors:** Laila Hugrass, Izelle Labuschagne, Eveline Mu, Ariane Price, David P Crewther

## Abstract

Functional imaging and behavioural studies have shown that the neuropeptide oxytocin influences processing of emotional faces. However, it is not clear whether these effects reflect modulation at an early or late stage of affective processing. We investigated the effects of oxytocin administration on early and late visual evoked potentials (VEP) in response to faces with neutral, fearful and happy expressions. In addition, we measured multifocal VEP and its associated nonlinearities to ascertain whether any changes observed in electrophysiology were indicative of a generalised effect or of one tied strictly to emotional processing. In a randomized, double-blind, cross-over design, 27 healthy male participants self-administered a nasal spray of either oxytocin (24 IU) or placebo. At very early latencies (40-60ms), oxytocin reduced right-temporal responses to fearful faces (*d* = .51), and central responses to both fearful (*d* = .48) and neutral faces (*d* = .54). For left occipito-temporal electrode sites, oxytocin decreased P100 reactivity to fearful expressions (*d* = 0.72). Oxytocin also decreased the amplitudes of the vertex positive potential (140-180ms) and late positive potential (400-600ms), regardless of whether the faces had fearful, happy or neutral expressions. The mfVEP showed no signs of selective magno-or parvo-cellular peak modulation comparing OXT with placebo with either low or high contrast stimulation. These results suggest that at early stages of visual processing, nasal oxytocin modulates responses to facial emotions, whereas at later stages of visual processing, it appears to influence more general face processing mechanisms. In addition, the measurable effects of OXT appear to be not a result of generalized brain change, but systematically related to emotional processing.

## Introduction

The neuropeptide oxytocin (OXT) is most commonly known for its powerful role in childbirth and mother-infant bonding, but has also shown to have a crucial role in modulating complex social behaviours in humans (Bartz and Hollander, 2008; Meyer-Lindenberg, 2008; Meyer-Lindenberg et al., 2011). Some authors have argued that this modulation can be traced to the effects of OXT on sensory processing of affective cues (Domes et al., 2007a). The amygdala plays an important role in processing social stimuli (Adolphs, 2008; LeDoux, 1998). Early amygdala responses to affective stimuli appear automatic in nature, not requiring conscious awareness, whereas later responses are modulated by attention, exhibiting tuning for behaviourally relevant input (Adolphs, 2008; Anderson et al., 2003). Functional imaging studies have shown that intranasal OXT administration can suppress amygdala responses to fearful and happy faces (Domes et al., 2007a; Kirsch et al., 2005), and can influence functional connectivity between superior colliculus and amygdala (Gamer et al., 2010) and frontal cortex (Sripada et al., 2012). Intranasal OXT has also been shown to improve recognition of facial emotions (Lischke et al., 2012; Marsh et al., 2010; Schulze et al., 2011). Interestingly, intranasal OXT normalizes amygdala responses to affective faces in groups with generalized social anxiety disorder and autism (Domes et al., 2013a; Domes et al., 2014; Labuschagne et al., 2010).

Due to the limited temporal resolution of fMRI, it is unclear whether OXT primarily influences affective processing at early or late stages of the visual hierarchy. Other techniques, such as visual evoked potentials (VEPs) have been applied to investigate the timing of affective processing (Hajcak et al., 2009; Pourtois et al., 2013). The visual P100 is a fast response that typically peaks between 80 and 120ms, and appears to originate from striate and extrastriate neural generators (Allison et al., 1999; Clark and Hillyard, 1996). Several studies have shown that P100 responses are influenced by the perceptual processing of facial expressions, with greater amplitudes in response to viewing fearful and happy faces than to neutral faces (Batty and Taylor, 2003; Vlamings et al., 2009). There is also evidence that viewing of facial emotions influences EEG and MEG activity even prior to the P100, as early as 40-60ms post stimulus presentation (Liu and Ioannides, 2010; Morel et al., 2012; Morel et al., 2009).

The N170 and the vertex positive potential (VPP) are responses that occur between 140 and 190ms, over right occipito-temporal and central sensors, respectively (Henson et al., 2003; Joyce and Rossion, 2005; Vlamings et al., 2009). Both the negativity and positivity reflect configural face processing, with greater responses to faces than objects, and to upright than inverted faces (Bentin et al., 1996; Joyce and Rossion, 2005). While these responses are sensitive to emotion, some evidence suggests that their amplitudes are modulated by emotional intensity, rather than valence (Luo et al., 2010). Although the literature has treated the N170 and VPP as two separate components, there is also evidence to suggest they reflect two sides of the same generator (Joyce and Rossion, 2005).

The late positive potential (LPP), a centrally generated, broad positive potential that begins 300 to 400ms after stimulus presentation (Crites et al., 1995), shows amplitude increase for both pleasant and unpleasant stimuli compared with neutral stimuli (Hajcak et al., 2009; Pastor et al., 2008). LPP amplitudes appear to reflect both automatic capture of attention, and goal-directed attention towards motivationally relevant stimuli (Hajcak et al., 2009).

There have not been many VEP studies into the effects of OXT administration on affective processing (reviewed, Wigton et al., 2015). In a sample of healthy females, Huffmeijer et al. (2013) investigated the effects of OXT administration on VPP and LPP responses to happy and disgusted faces, which were presented as performance cues in a flanker task. OXT increased VPP and LPP amplitudes, regardless of the emotional valence of the faces. However, the study did not include a neutral face condition, so it is unclear whether these results specifically reflect modulation of affective processing, or face processing mechanisms in general. Althaus et al. (2015) compared the effects of intranasal OXT on LPP responses to positive, negative and neutral scenes in males with functioning autism and a healthy male control group. There were no effects of OXT for either group; however, they found some effects of OXT on LPP responses to scenes featuring people, but only for healthy participants with high sensitivity to punishment, and for autistic participants with high blood plasma levels of OXT at baseline (Althaus et al., 2016). Differences in the effects of OXT on affective processing for these two studies (Althaus et al., 2015; Huffmeijer et al., 2013) may be due to the fact that OXT has a greater modulatory influence on amygdala responses to faces than to scenes (Kirsch et al., 2005).

To our knowledge, no previous studies have investigated the effects of OXT administration on the early visual responses to affective faces. We therefore compared early and late VEP responses to neutral, fearful and happy faces after the administration of nasal sprays containing OXT or a placebo. Previous studies have shown that OXT administration does not influence P100 or N170 responses to non-affective face stimuli (Herzmann et al., 2013; Rutherford et al., 2017). However, based on evidence that OXT administration inhibits amygdala reactivity to fearful and happy faces (Domes et al., 2007a), we hypothesized that it would reduce the effects of facial emotion processing on early (40-60ms), P100, N170 and VPP responses. Later potentials (LPP) are influenced by both stimulus salience and goal directed attention (Hajcak et al., 2009), and therefore it was unclear whether OXT would enhance or diminish the effects of facial emotion processing on LPP amplitudes. We also investigated whether individual differences in trait-level autism and social anxiety may influence the effects of OXT on VEP responses.

While the site and mechanism of action of nasal OXT remains to be fully settled, it is prudent to control for OXT having a general effect on the brain – eg through changing excitability, or through effects on functions such as blood pressure that might explain some of its effect on brain activity. Hence we measured the temporal structure of the nonlinear VEP, which is sensitive to the main afferent magnocellular and parvocellular visual streams (Klistorner et al., 1997), and to individual differences (Jackson et al., 2013; Sutherland and Crewther, 2010), and which is recorded using non-emotive flashed stimuli. Here we predicted that OXT would not have a measurable effect on the temporal structure of the simple VEP.

## Methods

### Participants

27 healthy males aged 18 to 40 (24 right handed, *M* = 25.22 years, *SD* = 4.72 years), gave written informed consent for the experiment, which was conducted with the approval of the Australian Catholic University Human Research Ethics Committee and in accordance with the code of ethics of the Declaration of Helsinki. One additional participant completed the placebo session but did not return for the oxytocin session and hence his results were excluded from the analysis. The happy facial expression condition was not added to the protocol until after the first four participants had completed the experiment, so we only have data for 23 participants in this condition.

### Questionnaires

Prior to attending the lab sessions, participants completed online versions of several scales. For the online emotion recognition test (Labuschagne et al., 2013), participants reported the emotion (anger, disgust, fear, happiness, sadness, and surprise) of 60 faces from the Ekman and Friesen face set (Ekman and Friesen, 1976). The Autism Spectrum Quotient (AQ) (Baron-Cohen et al., 2001) measures autistic personality traits, with higher scores indicating higher levels of trait autism (range, 0–50). The Social Interaction Anxiety Scale (SIAS) (Mattick and Clarke, 1998) was used to measure social anxiety, with higher scores indicating a greater degree of social anxiety (range, 0– 80). The State-Trait Anxiety Inventory (Spielberger, 2010) was administered to measure state anxiety prior to nasal spray administration (STAI_pre_), and the change in state anxiety by the end of the session (STAI_change_). State anxiety results from this sample are reported in more detail in Part 2 (Hugrass and Crewther, Submitted for Publication). In short, STAI_change_ was not significantly affected by OXT or PBO administration so this variable was redundant, and it was not included in the current analyses. We entered STAI_pre_ as a covariate in the preliminary models, to test whether subjective reports of state anxiety may influence the results.

### Facial emotion VEP task

The task was created using VPixx software (version 3.20, http://www.VPixx.com), and presented using a DATAPixx display driver and a Viewsonic LCD monitor (60Hz, 1024 x 768 pixel resolution) with linearized colour output (as measured with a ColorCal II – Cambridge Research Systems). Seven face identities (3 female) were selected from the Nimstim Face Set (Tottenham et al., 2009) with neutral, fearful and happy expressions (all with closed mouths, to minimise low-level differences between the stimuli). The images were converted to greyscale, the external features (hair, neck and ears) were removed, and the luminance and root mean square contrast were equated using a custom Matlab script (The Mathworks, Natick, MA). The stimuli were presented within a 20 × 19.5 degree mid-grey frame (47 cd/m^2^) on a grey background (65 cd/m^2^).

Participants were seated 70 cm in front of the display. A phase-scrambled neutral face (luminance and RMS contrast matched with the test stimuli) was presented during the baseline period (1800ms), followed by the target face (500ms), and a central fixation cross. After the face disappeared, participants used a RESPONSEPixx button box to report whether the expression was fearful (right button), neutral (top button) or happy (left button). Participants were told that this was not a speeded task and were instructed to respond as accurately as possible. In total, there were 240 trials (80 replications each, for the fearful, neutral and happy faces). To prevent fatigue, the task was divided into two blocks of 120 trials, with a self-timed break in between blocks.

### mfVEP Stimuli

The stimuli were presented on a 60Hz LCD monitor (ViewSonic, resolution 1024 x 768) with linearized colour output (measured with a ColorCal II). The 9-patch dartboard stimuli were created using VPixx software (version 3.20, http://www.VPixx.com), with a central patch (5.4° diameter) and two outer rings of four patches (21.2° and 48° diameter). The luminance for each patch was modulated at the video frame rate (60Hz, mean luminance = 42 cd/m ^2^, CIEx = 0.32, CIEy = 0.33), in pseudorandom binary M-sequences (M = 14), at either low (10% Michelson) or high (70% Michelson) temporal contrast. The M-sequences for each patch were maximally offset, so we could record independent responses across the visual field. For the purpose of this experiment, we only analyzed responses to the central patch.

### Procedure

Participants attended the laboratory twice, with at least a one-week washout period in between sessions. Participants were instructed not to drink alcohol on the night before their session, to refrain from drinking caffeine on the day of their session, and to refrain from eating or drinking (except for water) within an hour of their session. The order of treatment conditions was counterbalanced across participants. The spray bottles were relabelled so that neither the experimenters nor participants were aware of which spray bottle contained the oxytocin. Participants self-administered OXT (24 IU) or placebo sprays (PBO, containing all ingredients except for the peptide). Participants were given standardised instructions for how to administer the sprays, as per the recommended guidelines (Guastella et al., 2013). Spray bottles were first primed by puffing three sprays into the air, then participants inhaled a full spray in one nostril (4 IU) and waited 45s before inhaling a full spray into the opposite nostril. This was repeated until they had made three sprays per nostril. The facial emotion VEP task commenced at approximately 45 minutes after the last spray application (OXT: *M*= 43.81, *SD*= 4.00, PBO: *M* = 44.37, *SD*= 2.99).

### EEG recording and pre-processing

EEG was recorded using a 64-channel cap (Neuroscan, Compumedics). The data were sampled at 1KHz and band-pass filtered from 0.1-200Hz. The ground electrode was positioned at AFz and linked mastoid electrodes were used as a reference. EOG was monitored using electrodes attached above and below one eye. Data were processed using Brainstorm (Tadel et al., 2011), which is documented and freely available for download online under the GNU general public license (http://neuroimage.usc.edu/brainstorm).

### Emotional Face Processing ERP

EEG data were band-pass filtered (1- 40Hz), signal space projection was applied to remove eye-blink artefact and other segments of data containing low frequency noise (1 – 7Hz) were removed from the analysis. Baseline corrected (100ms to -1ms) epochs of data (−100 to 900ms) around the onset of each face presentation were imported into the Brainstorm database. Epochs containing high amplitude noise (>75μV) were excluded from the analysis. There were no consistent differences in the number of artefact contaminated epochs for the OXT and PBO session; however in order to minimise bias in each participant’s peak amplitude estimates (Luck, 2010), a subset of trials from the condition with more epochs was randomly selected so that the same number of trials contributed to the VEP waveforms for the OXT and PBO conditions (total number of epochs for the OXT and PBO sessions: Neutral = 1,942; Fearful = 1,948; Happy = 1,614).

P100, N170, VPP and LPP amplitudes were measured using LabVIEW (National Instruments), within time windows and electrode clusters that were based on previous research (Batty and Taylor, 2003; Luo et al., 2010; Pastor et al., 2008; Vlamings et al., 2009). Early responses were defined as the mean amplitude from 40-60ms post-presentation for the central (C1, Cz, C2, CP1, CPz, CP2, FC1, FCz, FC2), left (PO7, P7), and right (PO8, P8) electrode clusters (Liu and Ioannides, 2010). N170 and P100 responses were detected from right (PO8, P8) and left (PO7, P7) clusters of occipito-temporal electrodes. P100 was defined as the maximum amplitude between 80-120ms. N170 was defined as the minimum amplitude between 140-190ms. VPP was defined as the peak amplitude within the 140-190ms time-window for a central electrode cluster. In our analyses, the LPP response tended to be strongest from 400-600ms post-presentation, so LPP was defined as the average amplitude over the 400-600ms time window for the central cluster of electrodes.

### mfVEP

EEG data were band-pass filtered (1-40Hz) and signal space projection was applied to remove eye-blink artefact. Epochs of data were extracted from -100 to 500ms around the onset of each video frame (*n* = 16384). Each epoch was baseline corrected by subtracting the mean baseline amplitude (−100ms to -1ms). Custom Matlab/Brainstorm scripts were written for the mfVEP analyses in order to extract the K1, K2.1 and K2.2 kernel responses for the central patch. K1 is the difference between responses to the light and dark patches (Figure 1b). K2.1 measures neural recovery over one frame (16.67ms) by comparing responses when a transition did or did not occur (Figure 1c). K2.2 measures neural recovery over two frames (33ms), it is similar in form to K2.1, but includes an interleaving frame of either polarity (Figure 1d). Analyses of the contrast-response functions indicate that both the M and P pathways contribute to K1 amplitudes, K2.1 and early K2.2 responses originate from the M pathway, and that the main K2.2 response originates from the P pathway (Jackson et al., 2013; Klistorner et al., 1997).

**Figure 1:**
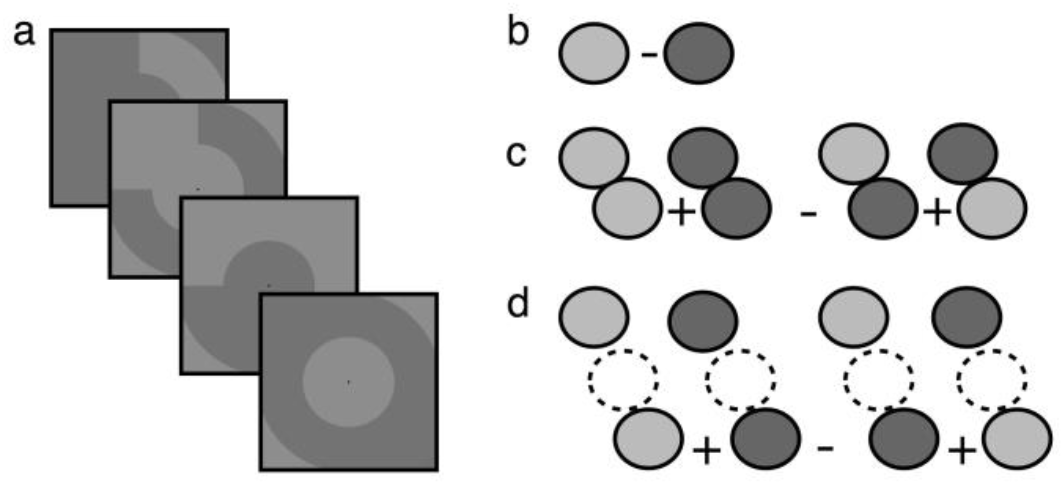
Illustration of the central two rings of the multifocal stimulus. (a) The patches alternate between light and dark grey in pseudorandom binary sequences, that are updated every video frame (60Hz). (b) The first-order kernel (K1) is the difference in summed response when the central patch was light or dark, K1=0.5*(S_L_-S_D_). (c) The first slice of the second order kernel (K2.1) takes the previous frame into consideration, comparing responses when a transition did or did not occur: K2.1=0.25*(S_LL_ + S_DD_ - S_LD_ - S_DL_). The second slice of the second order kernel (K2.2) is similar to K2.1, but there is another intervening frame of either polarity: K2.2=0.25*(S_L_L_ + S_D_D_ - S_L_D_ - S_D_L_).

LabVIEW (National Instruments) scripts were written to identify peak amplitudes and latencies in the VEP responses. Trough-to-peak K1_N65P105_, K2.1_N70P105_, K2.2_N70P90_ and K2.2 _N130P160_ amplitudes were exported for statistical comparisons. If a waveform could not be detected above the noise level, it was marked as a missing value (five out of 108 K2.1_N70P10_ responses were removed from the analysis).

### Statistical Analyses

The data were analysed using the linear mixed-effects modelling (LMM) procedure in SPSS, because of its advantage in dealing with missing values, and ability to handle unequal variance and correlated data. Separate analyses were performed for the behavioural reporting (accuracy and latency) dependent variables, and for each of the VEP dependent variables described above.

### Emotional Face Processing ERP

For the main analyses, important predictors were entered as covariates, and treatment condition (OXT vs. PBO), facial emotion (N, F, H) and the treatment by emotion interaction were entered as fixed effects. To account for individual differences in VEP amplitudes, random intercepts were modelled for each subject. In repeated-measures designs it is likely that the error terms are correlated within subjects. Hence, we used a first-order, autoregressive (AR1) covariance matrix for our models. The SPSS LMM procedure does not produce R^2^ effect sizes, because these definitions become problematic in models with multiple error terms. Where appropriate, Cohen’s *d* was calculated as the effect size for treatment and emotion comparisons.

For each result, we ran a preliminary LMM to investigate which of the variables contribute significantly to the models. For the analyses of behavioural response accuracy and latency, the preliminary covariates included AQ (*range =* 1-38, *M* = 17.48, *SD*= 7.71), SIAS (*range =* 11-50, *M* = 28.44, *SD*=1.69), and emotion recognition scores (*range =* 42-58, *M* = 51.87, *SD*= 3.82), as well as STAI_pre_ (as measured at prior to the nasal spray administration: OXT *range =* 20-58, *M* = 33.00, *SD*= 8.66, PBO *range =* 20-59, *M* = 32.22, *SD*= 9.60). To test whether other variables contributed to variation in behavioural responses, we also included time of day, and task latency (i.e., the time from the last nasal spray administration to the start of the facial emotion VEP task) as covariates. To take into account any relationship between accuracy and latency, response latency and accuracy were included in the respective models. Given that participants were instructed to wait to respond until after the face disappeared, we expect variation in response latencies to reflect the degree of difficulty in identifying the emotion, rather than the speed of identification per se.

For the VEP analyses, the preliminary LMMs included AQ, SIAS, STAI_pre_, behavioural response latency, time of day, and VEP task latency (from the time of the last spray administration) as covariates. To control for individual differences in behavioural reporting, we included response latency instead of response accuracy as a covariate, because these variables were correlated and there were ceiling effects for response accuracy. If a variable contributed significantly to the preliminary LMM, it was included as a covariate in the subsequent LMMs. For the P100 and N170 analyses, the fixed-effects for the LLM included factorial comparisons of the treatment, hemisphere, and emotion conditions. The VPP and LPP responses were recorded from a central cluster, so only treatment and emotion were included as repeated-measures fixed-effects in these analyses.

### mfVEP

In order to investigate the change in state anxiety over the testing session (STAI_change_), a preliminary mixed-effects analysis was conducted to investigate which variables should be included as predictors in the main model. Treatment was entered as the repeated-measures fixed effect and Subject ID was entered as the random intercept. AQ, SIAS and STAI_pre,_ order of treatment, time of treatment and latency of STAI_post_ were entered as covariates. Only STAI_pre_ contributed significantly to the preliminary model (see results section), so it was included as a covariate in the main LMM of STAI_change_.

Separate preliminary mixed-effects analyses were conducted to investigate important predictors of K1_N65P105_, K2.1_N70P105_, K2.2_N70P90_ and K2.2_N130P160_ amplitudes. The distributions of the K1_N65P105_, K2.1_N70P105_, K2.2_N70P90_ and K2.2 _N130P160_ amplitudes were all positively skewed. However, LMM does not require the DV to be normally distributed, and the model fits were similar with and without log_10_ transforms. Hence, in order to simplify interpretation of the estimated marginal means, analyses were performed on the untransformed amplitudes. For the preliminary analyses, treatment and contrast were entered as the repeated-measures fixed effects, AQ, SIAS, STAI_pre_, time of day, order of treatment and VEP latency were entered as covariates, and subject ID was entered as the random intercept. If any variables contributed significantly to the model for a given VEP waveform they were included in the main mixed-effects model for that waveform (see results section).

## Results

### Preliminary analyses of behavioural data

Preliminary LMMs were conducted on response accuracy and latency (as described above). Response latency (*F*(1, 69.3) = 7.47, *p* = .008), Response accuracy (*F*(1, 92.7) = 6.17, *p* = 0.015), and AQ (*F*(1, 16.7) = 14.31, *p* = 0.002), contributed significantly to the model of response latency, so these variables were included as covariates in the main analysis. Behavioural response latencies tended to be slower for participants with lower accuracies and with higher AQ scores (see Figure 1c and d). SIAS, STAI_pre_, emotion recognition scores, time of day, and VEP task latency did not contribute substantially to either model.

### Main analyses of behavioural data

Estimated marginal means (EMMs) for behavioural accuracy (corrected for response latency) and latency (corrected for response accuracy and AQ scores) are presented in Figure 2a and 2b. The face stimuli had high intensity expressions and were presented for 500ms, so it is not surprising that performance was often either at ceiling level or close to ceiling level. The mixed-effects analysis for response accuracy revealed a significant effect of emotion (*F*(2, 136.3) = 11.70, *p* < 0.001). Pairwise EMM comparisons (Bonferroni corrected) showed that accuracy tended to be higher for happy faces than for fearful (*p* = 0.028, *d* = 0.69) or neutral faces (*p* < 0.001, *d* = 1.09). The main analysis for response latency revealed a significant effect of emotion (*F*(2, 99.9) = 173.24, *p* < 0.001). Pairwise EMM comparisons (Bonferroni corrected) showed that latencies tended to be slower for neutral faces than for happy (*p* = 0.013, *d* = 0.43) or fearful faces (*p* < 0.001, *d* = 1.97). For both the accuracy and latency LMMs, there was no effect of treatment, and there was no interaction between the effects of treatment and emotion (*F* ≤ .33, *p* ≥ .57). In summary, regardless of treatment, responses tended to be more accurate for happy faces than for fearful and neutral faces, and slower for neutral faces than for affective faces.

**Figure 2:**
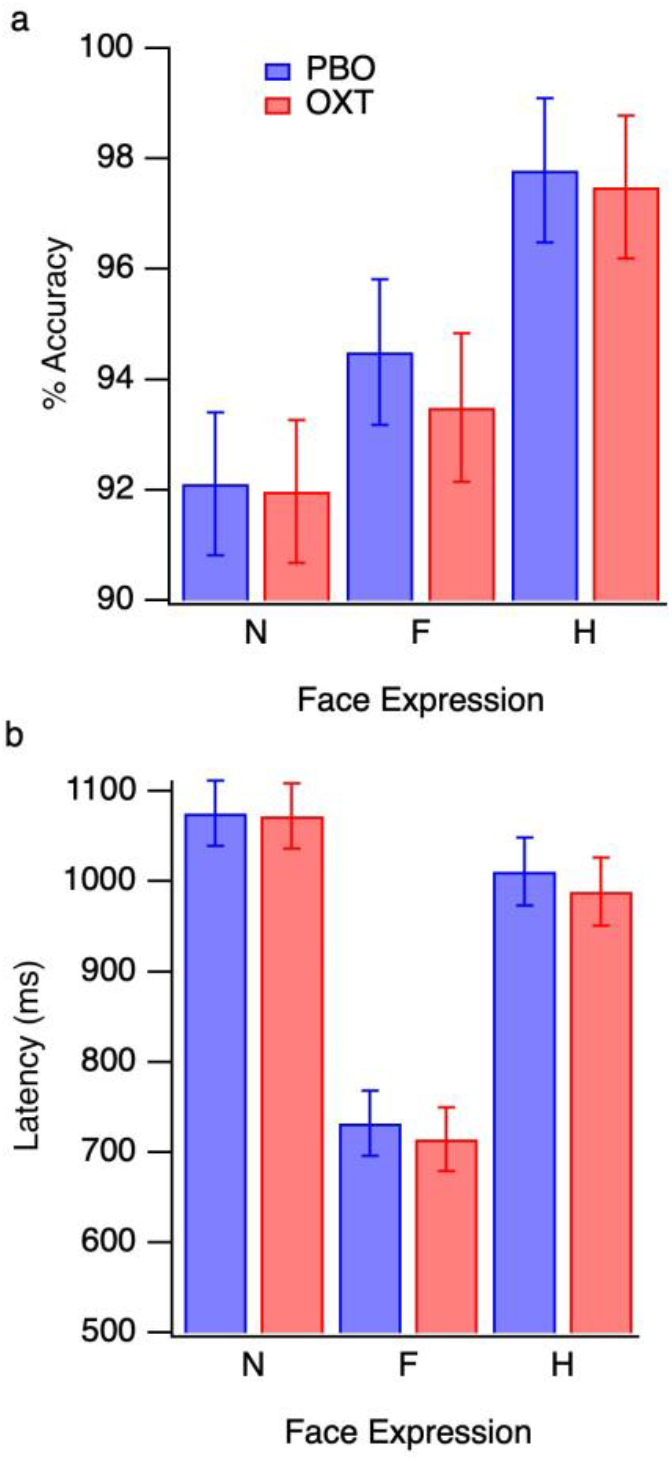
Behavioural results. Estimated marginal means for response accuracy (a) (corrected for individual differences in response latency), and (b) response latency (corrected for individual differences in response accuracy and AQ scores). Separate means are presented for the neutral (N), fearful (F) and happy (H) faces, error bars denote ±SEM. The results from the placebo (PBO) and oxytocin (OXT) sessions are presented in blue and red, respectively.

### Preliminary analyses of VEPs

Grand averages of the visual evoked potentials are presented in Figure 3, with separate traces for the neutral (*n* = 27), fearful (*n* = 27), and happy (*n* = 23) face conditions, and separate panels for the different electrode clusters and treatment conditions.

**Figure 3:**
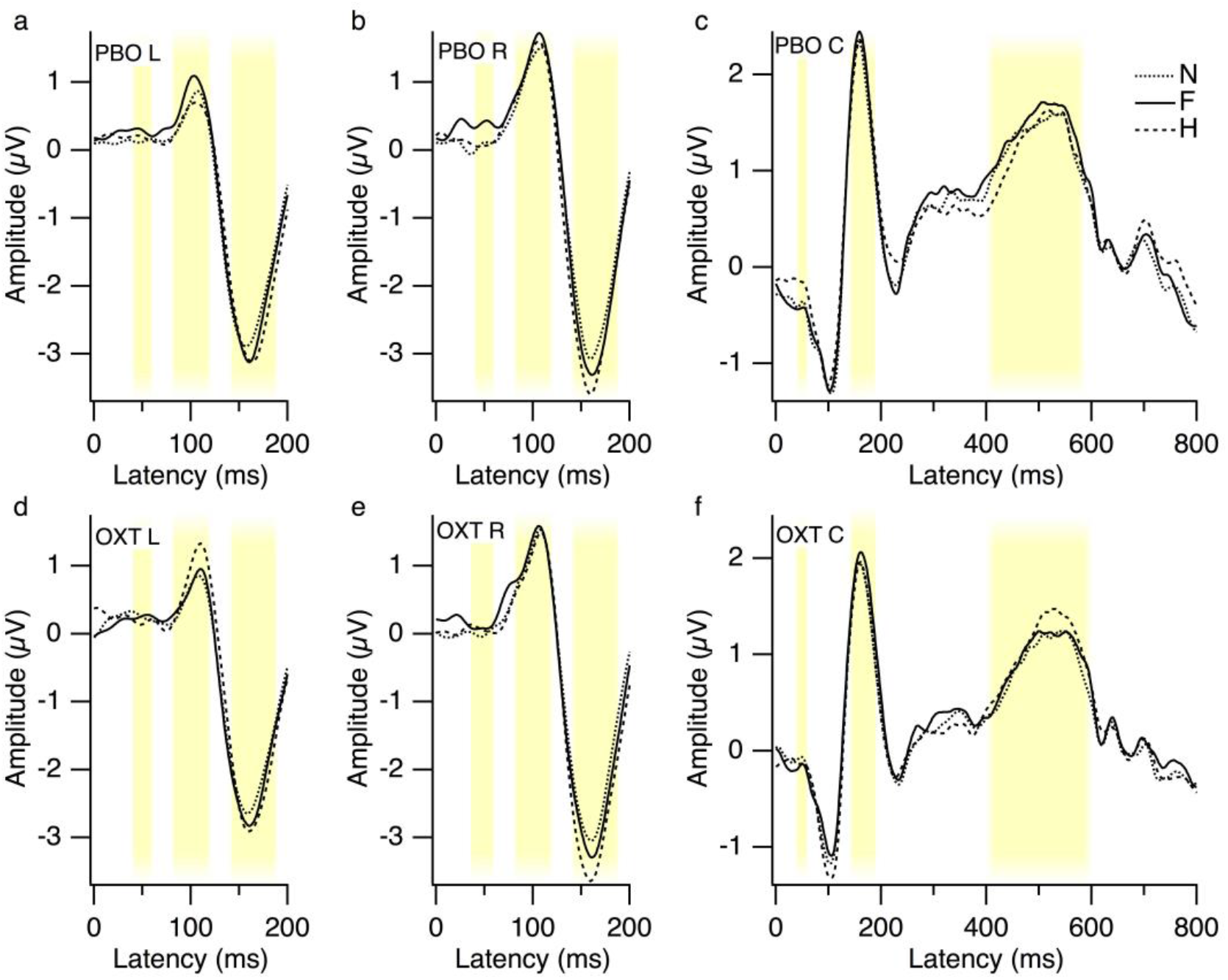
Grand mean visual evoked potentials. Panels (a), (b) and (c) present results from the placebo (PBO) session for the left (PO7, P7), right (PO8, P8), and central (C1-FC2) electrode clusters, respectively. Results from the oxytocin (OXT) session are presented for the left (d), right (e) and central (f) electrode clusters. Responses to the neutral (N), fearful (F) and happy (H) faces are presented in the dotted, solid and dashed traces, respectively. The yellow shading illustrates the time windows for the VEP analyses. Early responses from the clusters were averaged over the 40-60ms time window, P100 and N170 responses from the left and right clusters were detected within the 80-120ms and 140-190ms time windows respectively, whereas VPP and LPP responses from the central cluster were detected within the 140-190 and 400-600ms time windows respectively.

Preliminary LMMs (see Statistical Analyses section) of the early (40-60ms), P100, N170, VPP and LPP responses were conducted to identify which covariates to include in the main LMMs. Behavioural response latency contributed significantly to models of P100 amplitude, N170 amplitude, N170 latency, and LPP amplitude (*F* ≥ 5.00, *p* ≤.027), and the effect was approaching significance for early amplitude (40-60ms) (*F*(1,106.4) *=*3.67, *p* = .058) and P100 latency (*F*(1,145.2) *=*3.24, *p* = .074). At greater behavioural response latencies, VEP amplitudes tended to be smaller and VEP latencies tended to be longer. Hence, behavioural response latency was included as a covariate in the main LMMs for these dependent variables. SIAS contributed significantly to the model of early amplitudes for the central cluster (*F*(1,21.3) *=*5.32, *p* = .031); those with higher social anxiety tended to have stronger, central negativities during the 40-60ms time window. AQ contributed significantly to the preliminary LMM of N170 amplitudes (*F*(1,22.6) *=*8.92, *p* = .007), hence it was included as a covariate in the main N170 amplitude LMM. STAI_pre_, time of day and task latency did not contribute to any of the preliminary analyses, nor was there any evidence that they interacted with the effects of treatment, so these variables were not included in the main LMMs.

### Early effects (40-60ms)

We observed some very early effects of OXT administration. The VEP topographies and EMMs of amplitudes (averaged over the 40-60ms time window) are presented in Figure 4. We performed separate LMMs on these early amplitudes for the left, right and central electrode clusters. There were no effects of emotion or treatment on the left cluster amplitudes (*F* ≤ 1.04, *p* ≥ .36). The effect of treatment was significant for the central cluster (*F*(1, 44.9) = 8.99, *p* = 0.004), and the effect of emotion was approaching significance for the right cluster (*F*(2, 98.5) = 2.85, *p* = 0.063). Paired samples comparisons of the EMMs (Bonferroni corrected) revealed that for the right cluster, early responses to fearful faces were significantly reduced after OXT administration (*p* = 0.048, *d* = .51). For the central cluster, early responses to fearful (*p* = 0.013, *d* = .48) and neutral (*p* = 0.005, *d* = .54) faces were significantly reduced after OXT administration.

**Figure 4:**
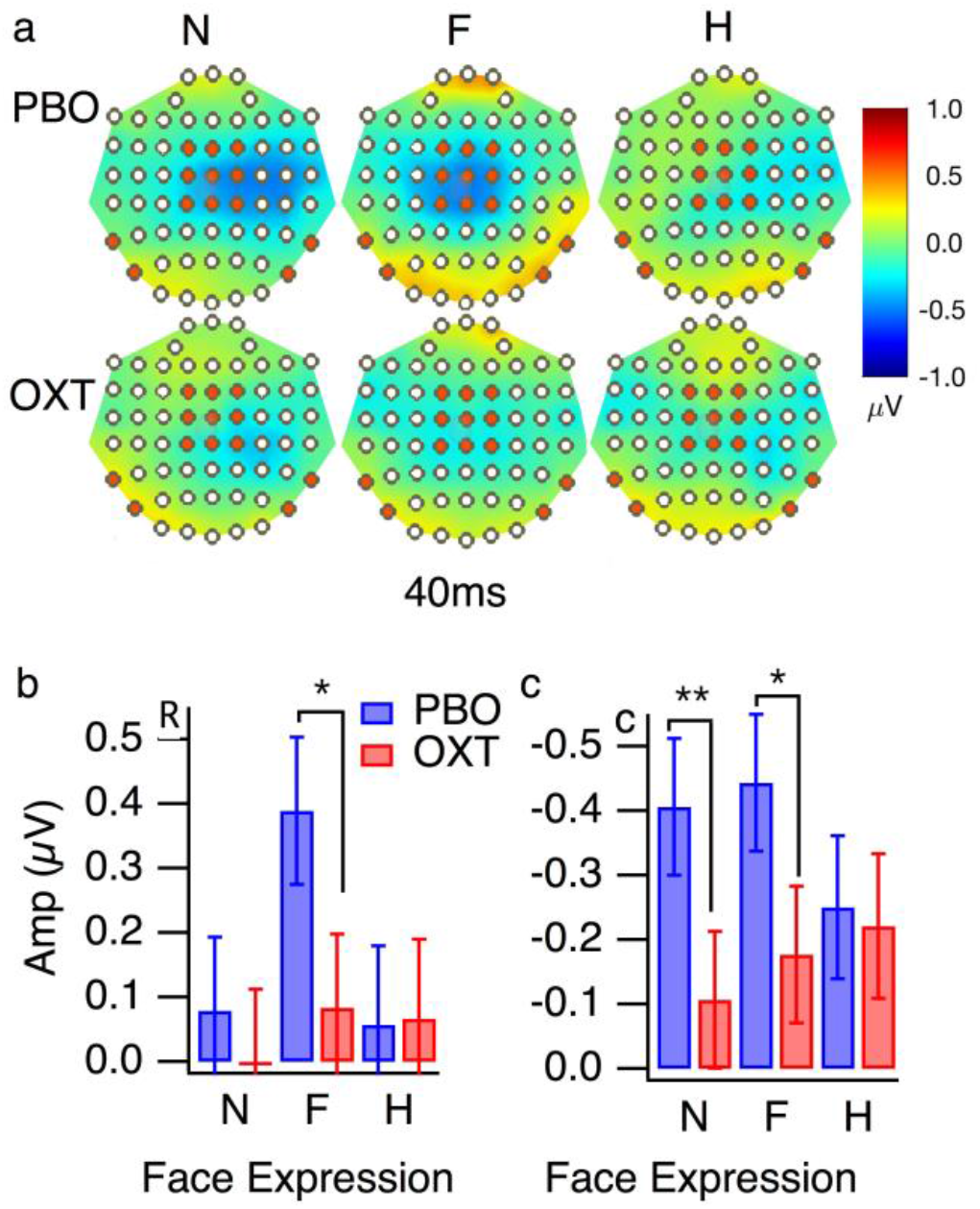
Very early VEP effects. (a) VEP topographies at 40ms for the neutral (N), fearful (F) and happy (H) faces, after placebo (PBO) and oxytocin (OXT) administration. The left, right and central electrode clusters are marked in orange. Panels (b) and (c) present the estimated marginal means amplitudes (averaged over the 40-60ms time-window) from the right and central electrode clusters, respectively. The y-axis for panel c is reversed, so that higher bars correspond with stronger negativities. Results from the PBO and OXT sessions are presented in the blue and red bars. The error bars denote ±SEM, * *p* < 0.05, ** *p* < 0.01.

Although the preliminary analyses showed that SIAS contributed significantly to the model for the central cluster, a follow-up analysis showed that it did not contribute significantly to the main analysis, nor was there any SIAS by treatment interaction (*F* ≤ 2.5, *p* ≥ .13).

### P100

EMMs for left and right P100 amplitudes and latencies (after correcting for behavioural response latencies) are presented in Figure 5. As illustrated in Figure 5b and c, the effects of emotion and treatment on P100 amplitudes were different for the left and right clusters. For the left cluster, there was a significant interaction between the effects of treatment and emotion (*F*(2, 98.5) = 14.48, *p* < 0.001). Pairwise EMM comparisons (Bonferroni corrected) showed that OXT decreased the amplitude of left P100 responses to fearful faces (*p* < 0.001, *d* = 0.72). There was also a trend towards OXT increasing the amplitude of left P100 responses to happy faces, but this effect did not reach statistical significance (*p* = 0.063, *d* = 0.43). However, the study was only sufficiently powered to detect strong effects, at least 64 participants would be required to detect weak to moderate treatment effects at 80% power (*d* =0.50; *α*=0.05). There were no other significant effects (*F* ≤ 1.69, *p* ≥ .19). For the right cluster, there were no effects of treatment or emotion and there was no emotion by treatment interaction (*F* ≤ 1.34, *p* ≥ .25). For LMMs of left and right P100 latencies (Figures 5d and 5e), there were no main effects or interactions for the emotion and treatment conditions (*F* ≤ 2.19, *p* ≥ .12).

**Figure 5:**
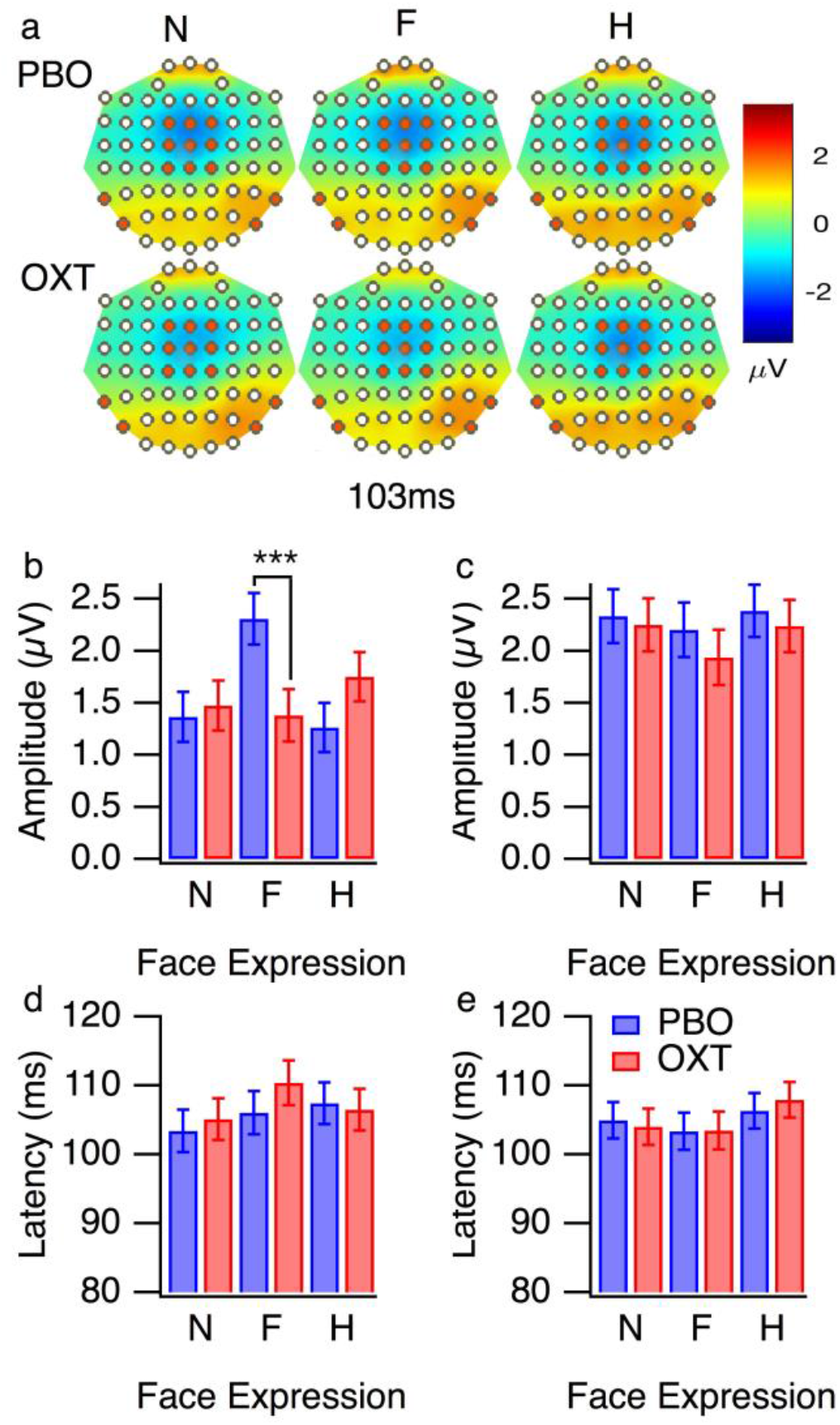
P100 results. (a) VEP topographies at 103ms for the neutral (N), fearful (F) and happy (H) faces, after placebo (PBO) and oxytocin (OXT) administration. The left, right and central electrode clusters are marked in orange. Estimated marginal means for (b) left P100 amplitude, (c) right P100 amplitude, (d) left P100 latency and (e) right P100 latency in response to neutral (N), fearful (F), and happy (H) faces. The estimated marginal means are corrected for individual differences in response latency. The results from the placebo (PBO) and oxytocin (OXT) sessions are presented in the blue and red bars. The error bars denote ±SEM, *** *p* < 0.001.

In summary, OXT administration modulates the effects of facial emotion on P100 amplitudes from electrodes over left occipito-temporal regions. It decreases P100 amplitudes for fearful faces and it may also increase P100 amplitudes for happy faces. However, it does not appear to influence right P100 amplitudes or P100 latencies from either hemisphere.

### N170 and VPP

N170 and VPP topographies are presented in Figure 6a, and EMMs for the amplitudes and latencies are presented in Figures 6b-g. Emotion contributed significantly to the LMM of right N170 amplitudes (*F*(2, 121.0) = 4.55, *p* = 0.012). Pairwise EMM comparisons (Bonferroni corrected) showed that right N170 amplitudes were greater (i.e., more negative) for happy faces than neutral faces (*p* = .010, *d* =0.26). The AQ and behavioural response latency covariates contributed significantly to the model of left N170 amplitudes (*F* ≥ 6.00, *p* ≤ .022), with greater amplitudes for longer behavioural response latencies and greater AQ scores. A follow-up analysis did not reveal any interactions between AQ and treatment. There were no other significant main effects or treatment by emotion interactions for the left or right N170 LMMs (*F* ≤ 1.55, *p* ≥ .22).

**Figure 6:**
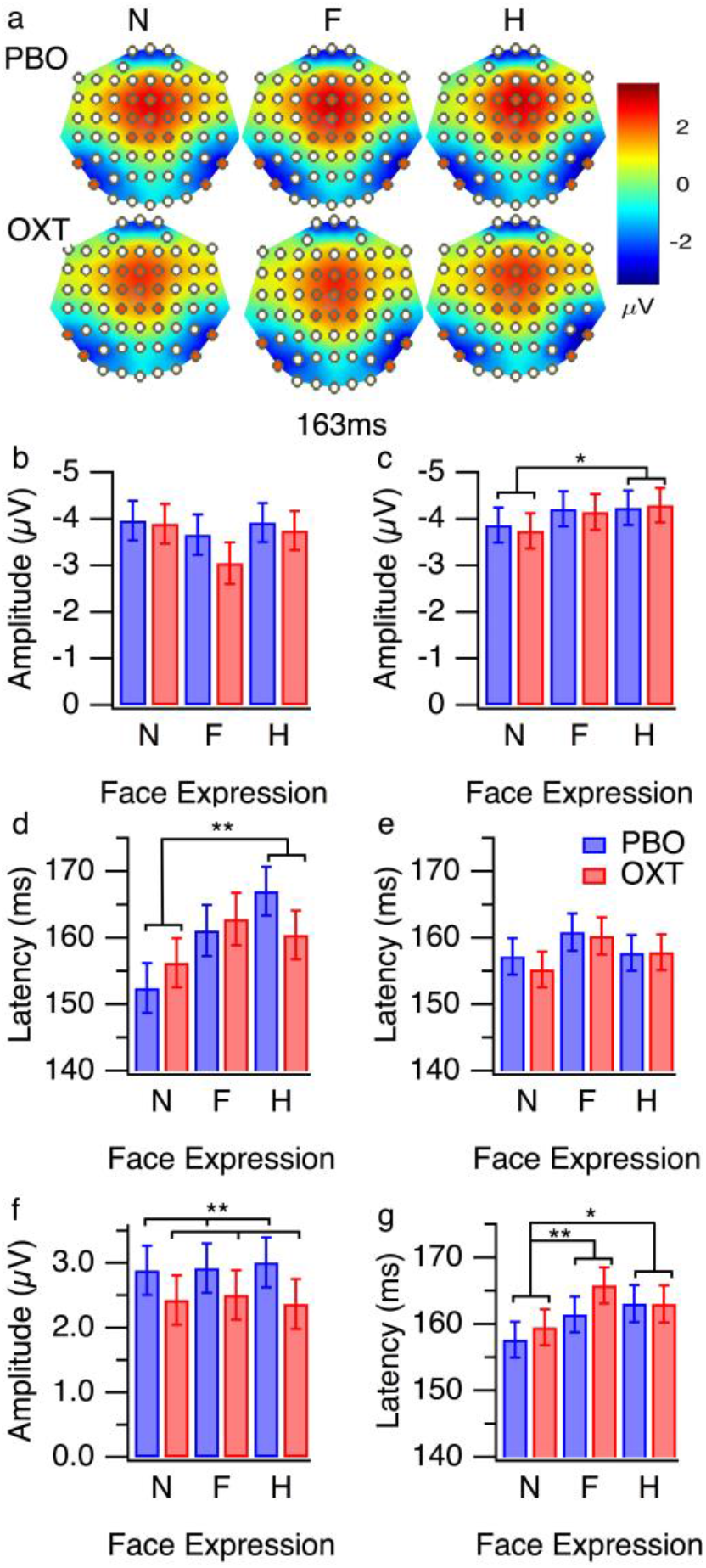
N170 and VPP results (a) VEP topographies at 163ms for the neutral (N), fearful (F) and happy (H) faces, after placebo (PBO) and oxytocin (OXT) administration. The left, right and central electrode clusters are marked in orange. Estimated marginal means (EMMs) are presented for (b) left N170 amplitude (c) right N170 amplitude, (d) left N170 latency, (e) right N170 latency and (f) central VPP amplitude (g) central VPP latency. The N170 EMMs (b-e) are corrected for individual differences in response latency. The N170 amplitude EMMs (b-c) are also corrected for individual differences in AQ scores. The y-axes for panels (b) and (c) are reversed, so that higher bars correspond with stronger negativities. The results from the placebo (PBO) and oxytocin (OXT) sessions are presented in the blue and red bars, respectively. The error bars denote ±SEM. * *p* < 0.05, ** *p* < 0.01

None of the effects for the right N170 latency LMM were significant (*F* ≤ 2.26, *p* ≥ .11). There was a significant effect of emotion on the left N170 latency LMM (*F*(2, 135.1) = 5.15, *p* = 0.007). Pairwise EMM comparisons (Bonferroni corrected) showed that left N170 latencies were longer for happy faces than neutral faces (*p* =0.005, *d* =0.64). There was no effect of treatment, and no treatment by emotion interaction (*F* ≤ 1.50, *p* ≥ .23).

For the model of VPP amplitude (Figure 6f), there was a significant effect of treatment (*F*(1, 36.5) = 7.40, *p* = 0.010, *d* =0.27), with reduced VPP amplitudes for OXT compared to PBO. However, there was no effect of emotion, nor was there an interaction between the effects of treatment and emotion (*F* ≤ .41, *p* ≥ .66).

For the model of VPP latency (Figure 6g), there was a significant effect of emotion (*F*(2, 93.8) = 5.93, *p* = 0.004). Pairwise EMM comparisons (Bonferroni corrected) showed that latencies were significantly shorter with neutral faces than fearful faces (*p* = 0.005, *d* =0.36) or happy faces (*p* = 0.034, *d* =0.31). However, there was no significant effect of treatment, nor was there a treatment by emotion interaction (*F* ≤ 1.55, *p* ≥ .22).

In summary, there were some effects of facial emotion on the N170 waveforms. Relative to neutral faces, happy faces produced higher right N170 amplitudes and slower left N170 latencies. However, regardless of AQ score, OXT did not influence N170 amplitudes or latencies. By contrast, VPP amplitudes tended to be reduced with OXT relative to PBO, regardless of facial emotion. VPP latencies tended to be shorter for neutral faces than affective faces, regardless of treatment.

### LPP results

LPP topographies and EMMs of LPP amplitudes are presented in Figure 7. Treatment contributed significantly to the LMM of LPP amplitude (*F*(1, 37.8) = 9.05, *p* = 0.005, *d* =2.22). This suggests that regardless of the facial emotion, LPP amplitudes to faces tend to be lower after OXT than after PBO. There was no significant effect of emotion, and no treatment by emotion interaction (*F* ≤ 1.85, *p* ≥ .16).

**Figure 7:**
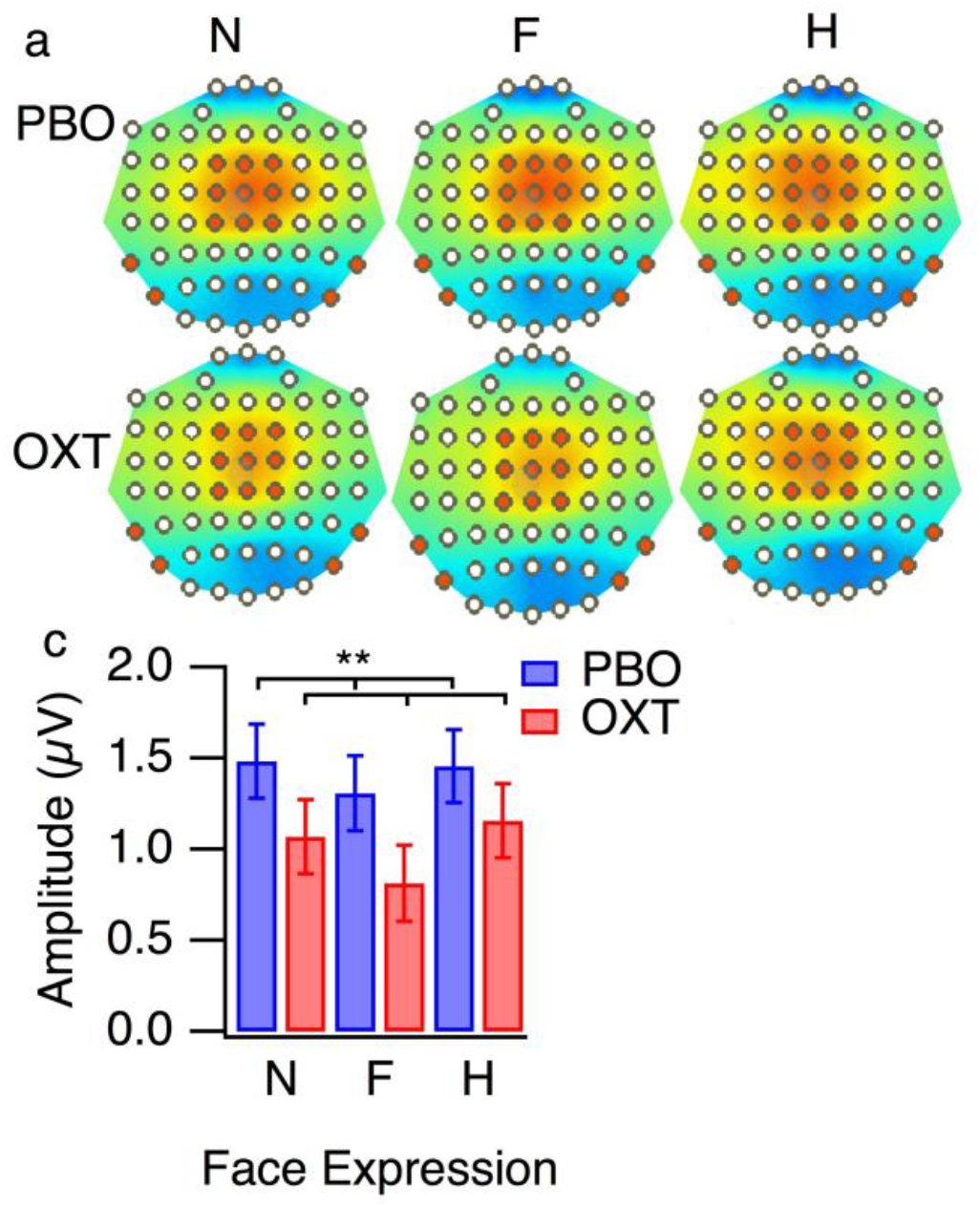
LPP results. (a) LPP topographies at 540ms for the neutral (N), fearful (F) and happy (H) faces, after placebo (PBO) and oxytocin (OXT) administration. (b) Estimated marginal means for LPP amplitude (corrected for behavioural response latency). The results from the placebo (PBO) and oxytocin (OXT) sessions are presented in the blue and red bars, respectively. The error bars denote ±SEM, ** *p* < 0.01.

### mfVEP Preliminary LMMs

The preliminary LMMs for K1_N65P105_, K2.1_N70P105_, K2.2_N70P90_ and K2.2_N130P160_ amplitudes revealed significant effects of SIAS on the K1_N65P105_ (*F*(1, 21.3) = 10.18, *p* = 0.004) and K21_N65P105_ *F*(1, 18.9) = 10.42, *p* = 0.004) amplitudes, so it was included as a covariate in the main LMMs for these waveforms. AQ, STAI_pre_, order of treatment, time of day and task latency did not contribute significantly to any of the preliminary mfVEP LMMs (*F* ≤ 2.66, *p* ≥ .12), so they were not included as covariates in the main analyses.

### K1_N65P105_ LMM

The K1 response (i.e.; the difference in response when the patch was light or dark) reflects the sum of inputs from the M and P afferent pathways (Klistorner et al., 1997). K1 waveforms for the low and high temporal contrast stimuli are presented in Figures 8a and 8d and estimated marginal means (EMMs) of K1_N65P105_ amplitudes (corrected for individual differences in SIAS) are presented in Figure 8g. The grand mean average waveforms show the expected increase in amplitude with contrast of the triphasic disturbance.

**Figure 8:**
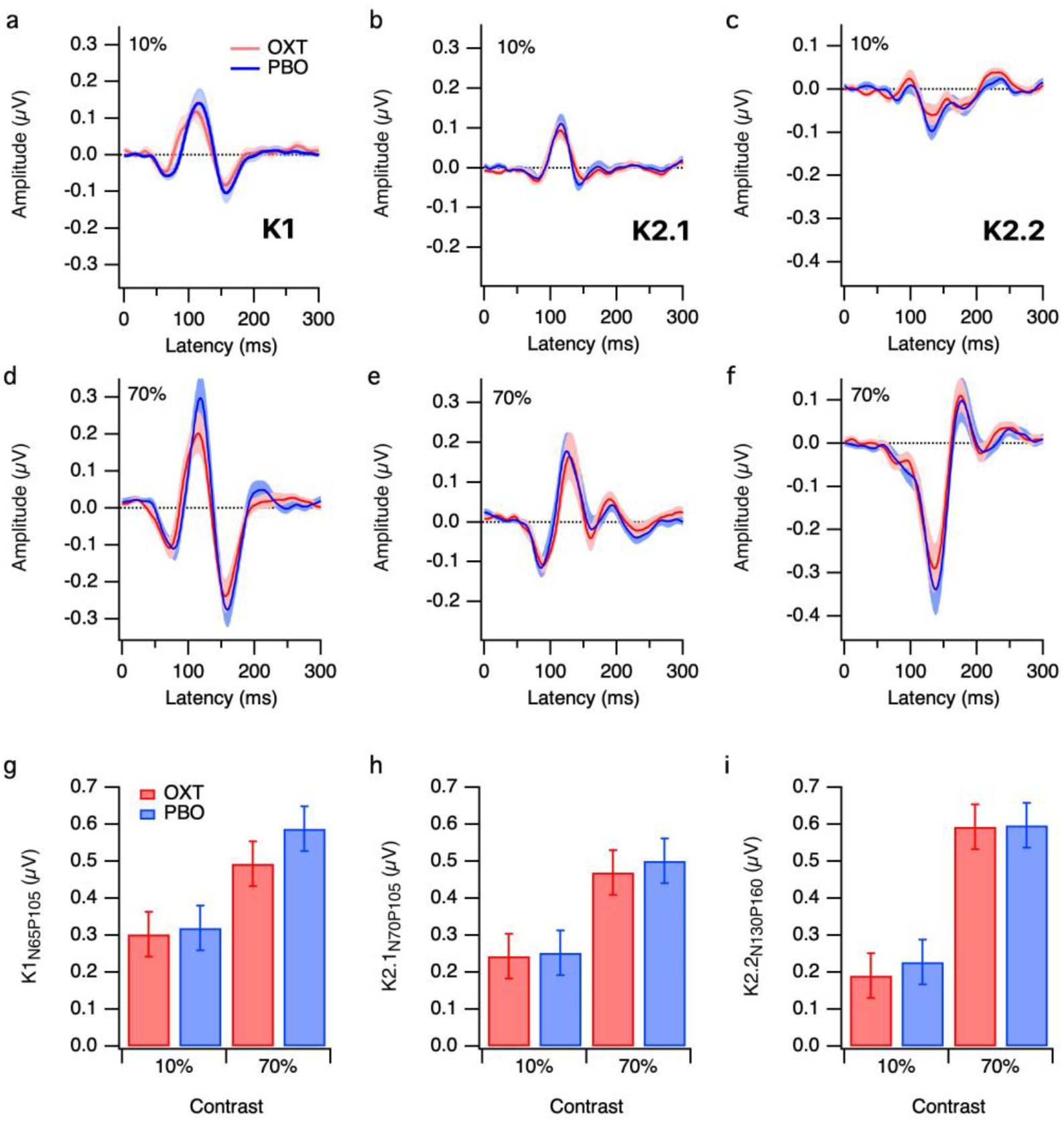
The effects of treatment (OXT and PBO) and contrast (10%, 70%) on (a) first order (K1), and (b, c) second order K2.1 and K2.2 slice responses, respectively. Mean waveforms for the low and high contrast stimuli are presented for OXT (red traces) and PBO (blue traces). The shading denotes ±1SE. Estimated marginal means of K1_N65P105_ amplitudes (g); K2.1_N70P105_ (h) ; K2.2_N130P160_ (i) (corrected for individual differences in SIAS) are presented for OXT (red bars) and PBO (blue bars) at low and high contrast.

An LMM on K1_N65_ latency (as measured at the peak negativity) showed an effect of contrast (*F*(1, 60.0)= 7.24, *p*= 0.010), but there were no treatment or treatment by contrast effects (*F* ≤ 2.48, *p* ≥ .12). There were no effects of contrast or treatment on K1_P105_ latency (as measured at the peak positivity), nor was there a contrast by treatment interaction (*F* ≤ 2.45, *p* ≥ .12). There appears to be a small advance in K1 latencies under OXT administration (Figures 8a) at low contrast. Similarly, the averaged differential (OXT-PBO) waveform at low contrast, for early latencies (20-70ms) demonstrates a systematic deviation from zero. This suggests that if OXT administration influences K1 latencies, it does not do so at the major peaks and the effects are small in magnitude.

The main mixed-effects analysis revealed there were significant effects of contrast (*F*(1, 48.7)= 41.50, *p*< 0.001) and SIAS (*F*(1, 24.9)= 7.43, *p*= 0.012) on K1_N65P105_ amplitudes. However, there was no significant effect of treatment, no treatment by contrast interaction and no SIAS by treatment interaction (*F* ≤ .98, *p* ≥ .33). Bivariate correlations showed that K1_N65P105_ amplitudes tended to be greater for people with higher SIAS scores, both for the low (OXT *r* = .39, *p* = .045; PBO *r* = .45, *p* = .017) and high contrast stimuli (OXT *r* = .39, *p* = .046; PBO *r* = .44, *p* = .022). In summary, our results suggest that K1_N65P105_ amplitudes tend to increase with social anxiety and stimulus contrast, but they are not affected by OXT vs. PBO.

### K2.1_N70P105_

The K2.1 response (i.e.; recovery of flash VEP over one video frame) is likely to originate from M inputs (Klistorner et al., 1997). K2.1 waveforms for the low and high temporal contrast stimuli are presented in Figures 4a and 4b, and EMMs of K2.1_N70P105_ amplitudes (corrected for individual differences in SIAS) are presented in Figure 4c. The grand mean waveforms show the expected increase in amplitude with contrast.

The LMM of K2.1 _N70P105_ amplitudes revealed significant effects of contrast (*F*(1, 29.5) =32.89, *p*<0.001) and SIAS (*F*(1, 24.5) =7.16, *p*= 0.013) on K2.1_N70P105_ amplitudes. However, there was no significant effect of treatment and there was no treatment by contrast interaction (*F* ≤ 1.23, *p* ≥ .27). Bivariate correlations (Figure 4d and 4e) showed that K2.1_N70P105_ amplitudes tended to be greater for people with higher SIAS scores for the PBO session (10% contrast *r* = .54, *p* = .005; 70% contrast *r* = .41, *p* = .043), but these correlations were not statistically significant for the OXT session (10% contrast *r* = .36, *p* = .060; 70% contrast *r* = .26, *p* = .196). However, the LMM showed that the SIAS by treatment interaction was not statistically significant (*F* =2.07, *p* = .16). Consistent with the analysis of K1 amplitudes, our results suggest that K2.1 amplitudes increase with stimulus contrast and social anxiety, but they are not substantially affected by OXT vs. PBO administration.

### Early and late K2.2 waveforms

The early and late K2.2 responses (i.e.; recovery of flash VEP over two video frames) are likely to originate from M and P inputs respectively (Jackson et al., 2013; Klistorner et al., 1997). The averaged K2.2 waveforms for the low and high contrast conditions are presented in Figures 8c and 8f respectively and EMMs of the late K2.2 amplitude is presented in Figures 8i. The grand mean waveforms show the expected increase in amplitude with contrast for the late K2.2 waveform. SIAS did not contribute significantly to the early or late K2.2 LMMs, so it was not included as a covariate for these analyses. The early, K2.2 _N70P90_ amplitude is small and affected by the development of the N130 at high contrast, and hence will not be discussed further.

The LMM of the late K2.2 _N130P160_ amplitudes revealed a significant effect of stimulus contrast (*F*(1, 24.5)=31.19, *p*<0.001). There was no effect of treatment, nor was there a treatment by contrast interaction (*F* ≤ .41, *p* ≥ .53) Our results suggest that neither early nor late K2.2 waveforms are affected by OXT vs. PBO administration.

## Discussion

### Emotional Face Processing ERP

To our knowledge, this is the first study to report modulation of early VEP responses to facial emotion after OXT administration. We found that OXT administration diminished the effects of facial emotion on early VEP amplitudes. After PBO treatment, central VEP amplitudes from 40-60ms discriminated between happy expressions and fearful or neutral expressions, whereas early right (40-60ms) and later left P100 amplitudes discriminated between fearful expressions and neutral or happy expressions. The latencies and topographies of these early effects are broadly consistent with the existing EEG and MEG literature (Liu and Ioannides, 2010; Morel et al., 2012; Morel et al., 2009). Affective discrimination at these early latencies is unlikely to rely on the geniculo-cortical visual processing route; rather it is thought to reflect a fast subcortical route to the amygdala, via the superior colliculus and pulvinar (Liddell et al., 2005; Vuilleumier et al., 2001). The decrease in early VEP responses to facial affect after OXT administration are interesting in light of psychophysical evidence that OXT improves recognition of briefly (18-53ms) presented happy and angry faces (Schulze et al., 2011). Taken together, these results support the theory that OXT modulates the salience of social cues at very early, automatic stages of affective processing (Bartz et al., 2011; Ebitz et al., 2013).

The effects of OXT on left posterior P100 amplitudes depended on facial emotion, with reduced amplitudes for fearful faces and a trend towards increased amplitudes for happy faces. Consistent with previous studies that only used non-affective face stimuli (Herzmann et al., 2013), OXT did not influence P100 responses to faces with neutral expressions. We did not observe any effects of OXT on P100 amplitudes from right, posterior electrodes. This is surprising because affective processing of faces tends to be biased towards the right hemisphere (Ley and Bryden, 1979). Studies have shown that different aspects of affective processing involve the left (Morris et al., 1998; Vuilleumier et al., 2001) or right amygdala (Adolphs et al., 1996; Pegna et al., 2005). Likewise, acute OXT administration can modulate right (Domes et al., 2007a), left (Domes et al., 2014) or bilateral (Labuschagne et al., 2010) amygdala reactivity to emotional faces. The neuromodulatory role of OXT on face processing appears to be more complicated than one would predict based on a simple right-hemisphere dominated model of emotional processing.

Despite the fact that N170 and VPP responses occur at the same time, and may share a common generator (Joyce and Rossion, 2005), we observed differences in the effects of OXT administration on the two potentials. Across the facial emotion conditions, VPP amplitudes tended to decrease after OXT administration, whereas N170 amplitudes were unaffected by treatment condition. Huffmeijer et al. (2013) found that OXT increased the amplitude of VPP responses to disgusted and happy faces. They interpreted this finding as evidence that OXT modulates early affective processing, but their experiment did not include a neutral face condition. Our results indicate that the effects of OXT administration on VPP responses are not specific to affective stimuli, and may reflect a more general modulation of face processing. However, even if this is the case, it is unclear why OXT modulated VPP but not N170 responses.

Consistent with our VPP analyses, OXT administration decreased the amplitude of LPP responses to faces with neutral, fearful and happy expressions. This is contrary to Huffmeijer et al. (2013), who found that OXT increased the amplitude of VPP and LPP responses to disgusted and happy faces. It is important to note that Huffmeijer et al.’s participants were healthy females and ours were healthy males. There are significant sex differences in brain responses to OXT; fMRI studies have found that OXT suppresses amygdala reactivity to affective faces in healthy males (Domes et al., 2007a) while enhancing it in healthy females (mid-luteal phase) (Domes et al., 2010). Future VEP studies that directly compare the timing of affective processing in male and female participants are required to elucidate potential sex differences in the neuromodulatory effects of OXT on affective processing.

We did not observe any effects of OXT administration on the accuracy or latency of emotion identification. In order to evoke strong electrophysiological responses, our face stimuli had strong emotional intensity, and long presentation durations (500ms). This meant that identification accuracy was close to ceiling level in both the PBO and OXT conditions. Previous studies have shown that OXT has greater effects on emotion identification for face stimuli with low intensity expressions (Lischke et al., 2012; Marsh et al., 2010) and short presentation durations (i.e., 100ms *cf*. 500ms) (Domes et al., 2013b). Hence, our results are not necessarily inconsistent with the literature on the effects of OXT on facial emotion identification.

It is well known that there are individual differences in the effects of OXT on social processing (Bartz et al., 2011). Previous studies have shown that OXT has differential effects on neural responses to emotional faces in groups with autism and generalized social anxiety disorder, relative to healthy control groups (Domes et al., 2013a; Labuschagne et al., 2010, 2012). Although there were wide ranges of AQ and SIAS scores in our sample (see analysis section), we did not specifically recruit people with extreme scores. Given the small sample size, our study was not sufficiently powerful to detect weak to moderate correlations, if they exist at the population level.

### mfVEP

In summary, OXT had no effect on state anxiety, or on any of the nonlinear VEP kernel amplitudes. Furthermore, there were no OXT by stimulus contrast interactions. Based on these results, there is no evidence that OXT modulates peak activation of the primary visual cortex via either the M or P afferent pathways (when using simple flash stimuli). As expected, based on the existing literature, kernel responses were greater in amplitude for the high contrast stimulus, and the effects of contrast on amplitude were stronger for the P-driven (late K2.2) responses than for the M-driven (K2.1 and early K2.2) responses (Jackson et al., 2013; Klistorner et al., 1997). Interestingly, some of the individual variation in VEP amplitudes could be attributed to the participants’ scores on a self-reported social anxiety measure. Specifically, M-driven (K2.1) amplitudes increased with social anxiety, whereas P-driven (late K2.2) amplitudes were not significantly associated with social anxiety. By contrast, autistic tendency did not predict non-linear VEP amplitudes. We provide more detailed interpretations of the effects of OXT, anxiety and autistic tendency on visual processing below.

There are several possible explanations as to why OXT did not affect VEP amplitudes. Differences in OXT spray administration methods can influence its dosage and bioavailability (Bradfield, 1965). Although we did not measure blood or salivary levels of OXT, we followed procedures to maximise the likelihood of the drug being absorbed efficiently at the mucosal surface (Guastella et al., 2013). The latencies from nasal spray delivery to the VEP and STAI measurements were well within the periods for which salivary OXT remains elevated (van Ijzendoorn et al., 2012), and peripherally measured OXT concentrations tend to be reliable indicators of central OXT concentrations after nasal spray administration (Valstad et al., 2017). Although the sample size was relatively small, many studies have reported effects of OXT on brain activation for crossover designs with similar or smaller sample sizes (Domes et al., 2007b; Kirsch et al., 2005; Labuschagne et al., 2010; Perry et al., 2010; Sripada et al., 2012).

The effects of OXT can vary depending on the context in which it is administered (Bartz et al., 2011). According to the social salience hypothesis, OXT enhances the salience of social cues (Ross and Young, 2009; Schulze et al., 2011). There is evidence that the oxytocin receptor (OXT-R) gene impacts functional activation in the visual cortex (O’Connell et al., 2012). Furthermore, OXT enhances functional coupling between the amygdala and the superior colliculus (Gamer et al., 2010), a subcortical region involved in gaze direction, which receives predominantly M input (Wurtz and Albano, 1980). While it is plausible that OXT could modulate processing in the afferent pathways, the results presented here and in Part 1 (Hugrass et al., Submitted for Publication) indicate that it only modulates early visual processing when there is socially salient input.

In addition to contextual effects, stable individual differences, such as endogenous plasma OXT levels and personality variables can also influence the effects of OXT administration on brain and behavioural measures (Bartz et al., 2011). Previous studies have shown that OXT has greater effects on amygdala activation in generalised social anxiety disorder and in autism than in control groups (Domes et al., 2013a; Labuschagne et al., 2010). Trait anxiety can also moderate the effects of OXT on brain and behavioural measures in non-clinical samples (Alvares et al., 2012). Therefore, we were interested in whether social anxiety levels in a neurotypical sample contribute to individual variation in non-linear VEP amplitudes. High social anxiety was associated with higher amplitudes for the M-driven (K2.1) nonlinear VEP amplitudes. This is consistent with evidence that trait anxiety improves M processing of ambiguous threat cues (Im et al., 2017). Correlations between K2.1 amplitudes and social anxiety were weaker after OXT administration; however our study was not sufficiently powerful to detect whether OXT blunts the effect of SIAS on M-driven VEP amplitudes at the population level.

We did not observe any effects of autistic tendency on M or P driven VEP amplitudes. By contrast, previous studies have shown that M-driven K2.1 amplitudes tend to be higher in neurotypical groups with high, compared with low, autistic tendency (Burt et al., 2017; Jackson et al., 2013). It is important to note that these studies were specifically designed to compare groups with low and high AQ scores, whereas we simply recorded AQ in order to control for any potential interactions between the effects of AQ and OXT administration on VEP amplitudes. Although our sample included some participants with very low and high AQ scores, the majority scored in the mid-range. Previous studies have shown that OXT administration can improve face processing in groups with ASD (Domes et al., 2007a), and furthermore that M abnormalities may contribute to differences in affective face processing in groups with low and high AQ scores (Burt et al., 2017). Therefore, it is reasonable to ask whether OXT administration can reduce the effects of autistic tendency on M-driven VEP amplitudes. In order to answer this question, future studies will require larger numbers of participants at the low and high ends of the autistic personality spectrum.

In summary, we investigated differences in the acute effects of nasal OXT administration on the non-linear temporal structure of visual evoked potentials. We were specifically interested in whether OXT would increase temporal efficiency in the M pathway; however, the absence of any treatment or treatment by temporal contrast effects on non-linear VEP amplitudes suggest that OXT does not influence processing in the main afferent streams to the primary visual cortex. High levels of social anxiety were associated with increased M-driven non-linear VEP amplitudes; this suggests that anxiety reduces temporal efficiency in the M pathway. However, this effect was not significantly different for the sessions when participants received OXT or PBO. Therefore although nasal OXT has been shown to bias top-down attentional resources towards socially relevant visual stimuli (Bartz et al., 2011), our results indicate that it does not play a role in gating non-social M and P input to the primary visual cortex.

## Conclusions

In summary, analysing VEPs enabled us to study the effects of OXT administration on the early and late stages of affective processing. OXT administration decreased reactivity to fearful expressions as early as 40-60ms for central and right electrode clusters, and as early as the P100 for left electrode clusters. It may also increase left P100 reactivity to happy expressions. By contrast, OXT administration decreased VPP and LPP amplitudes regardless of facial emotion. Our results indicate that OXT modulates the salience of social cues at very early stages of visual processing (Bartz et al., 2011; Ebitz et al., 2013), and that it modulates more general face processing mechanisms at later stages of visual processing. Importantly, early visual cortical changes under OXT administration, appear to relate only to affective stimuli, and not to the general mfVEP which uses flashes of simple diffuse stimuli. This study thus confirms that nasal OXT affects the electrophysiological response of the brain to emotional stimuli, but that the temporal structure of the nonlinear VEP appears unaltered by OXT, ruling out systemic or generalized brain activation as the cause.

## Acknowledgements

We are grateful to the undergraduate students who helped to collect the data. LH and DC were supported by the Australian Research Council (ARC) (150104172). Nasal sprays were purchased with funding from Swinburne University of Technology and the Australian Catholic University.

## Notes

### Competing Interest Statement

The authors have declared no competing interest.

### Summary of Updates

The text is unaltered. We unfortunately left out one of the agreed authors during submission, Eveline Mu, who played a critical role in improving and carrying out EEG analyses for the paper as well as reviewing the manuscript and its discussion.

